# Information neutrality generates robust biological scaling in stochastic systems

**DOI:** 10.64898/2026.01.19.700411

**Authors:** Andrea Tabi

## Abstract

Universal scaling laws are a signature of complex biological systems, yet the mechanisms by which microscopic stochastic processes generate robust macroscopic behaviour remain debated. Here we develop an information-theoretic approach to identify robust biological scaling using a stochastic ontogenetic growth model, in which correlated cellular bursts propagate from microscopic noise to organism-level metabolic fluctuations. We show that an information-neutral regime, defined as the minimum Kullback–Leibler divergence between fluctuation distributions generated under different microscopic correlation structures, identifies stochastic regimes for which macroscopic observables become maximally insensitive to microscopic details. Within this regime, metabolic scaling exponents converge across species sizes to a narrow range close to linear scaling relationship. The information-theoretic optimum is robust to temporal coarse-graining, indicating non-critical scale robustness without requiring proximity to a phase transition. Our results suggest that information neutrality provides a general principle for identifying robust stochastic mechanisms underlying biological scaling and illustrate how universal macroscopic behaviour can emerge from noisy microscopic dynamics.

## I. INTRODUCTION

Power-law distributions are prevalent in natural complex systems. This distribution is often interpreted as a signature of phase transitions which emerge when a control parameter is tuned to a critical threshold [29]. This underlies the concept of self-organized criticality, i.e. when natural complex systems evolve towards a critical threshold in order to achieve maximal information processing [16, 22]. Since “life at the edge of chaos” argument has gained traction in biology over the past decades, many biological processes have been observed to reside close to phase transitions [14, 30]. However, various mechanisms can generate power laws without criticality [37], such as aggregation [25], preferential attachment [4], or stochastic processes [31, 39]. Thus identifying the generative process remains crucial for understanding the biological significance and interpretation of such scaling laws.

Scaling laws are widespread across biological hierarchies. Traits including metabolic rate, growth, lifespan, and population density exhibit systematic relationships with organism size [3, 6, 35, 42]. These regularities suggest that biological systems obey underlying constraints governing how resources, fluctuations, and information propagate across organizational levels. Among these relationships, metabolic scaling is one of the most extensively studied examples of biological scaling laws, which describes the sublinear relationship between body size and metabolic rate rooted in an empirical observation called the Kleiber’s law [5, 19]. Since body size is a master trait, metabolic scaling belongs to a broader class of biological allometries [35] with life-history traits, population growth rates, and energy fluxes [7] to name a few. This suggests that living systems obey internal constraints as a consequence of historical contingencies and functional and structural rules [1, 38]. The universality of this relationship has been up for debate for a long time due to largely varying observations [8, 10, 12, 17], however the most popular explanations include non-critical phenomena such as fractal-like structures of distribution networks [41], surface-to-volume ratio of the organisms [15, 34] or life-history optimisation [43]. In biological contexts, such scale invariance may reflect constraints on how fluctuations and information propagate across biological hierarchy [40]. Coarse-grained variables such as metabolic rate are therefore informative macroscopic descriptors that captures the system behavior as a whole. Information theory provides a natural framework for quantifying how microscopic fluctuations influence macroscopic observables and how information about microscopic parameters is preserved or lost under coarse-graining [9, 40]. In complex systems, coarse-graining often leads to regimes in which macroscopic observables become insensitive to microscopic details, a property associated with universality in statistical physics [37, 44]. Recent work shows that metabolic fluctuations can successfully reproduce the observed variability of the metabolic scaling exponents across and within taxa [39]. Biological systems are inherently stochastic that may arise from various cellular processes such as birth–death dynamics, metabolic bursts due to thermodynamic fluctuations, noisy biochemical reactions, and stochastic gene expression [11, 26, 33, 36]. Such microscopic fluctuations can propagate across organizational levels, generating macroscopic variability even without environmental forcing.

Similarly, demographic noise in ecological systems arises from discrete birth-death processes, and is a fundamental source of intrinsic stochasticity [21, 32], which can generate macroscopic population fluctuations even without environmental forcings [27]. This raises a question under what conditions do macroscopic patterns, in this case the allometric exponents, become universal given the underlying microscopic noise structure.

In this study we investigate whether universal metabolic scaling can emerge from statistical properties of stochastic cellular dynamics. We present a stochastic ontogenetic growth model in which cellular bursts generate metabolic fluctuations whose magnitude scales with organism size. Using an information-theoretic analysis, we identify a regime where organism-level fluctuation statistics become minimally sensitive to changes in the microscopic correlation structure, i.e., an information-neutral regime. At this point, the macroscopic metabolic scaling exponents converge across species sizes. These results suggest that robust biological scaling laws may arise from information-theoretic properties of stochastic cellular processes rather than from criticality.

## II. METHODS

### A. Stochastic model of adult metabolism

We consider an adult organism composed of *N* cells.

Its body mass is approximated as

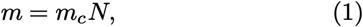

where *m*_*c*_ is the mean cellular mass. The deterministic maintenance metabolism is assumed to be proportional to cell number,

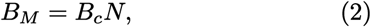

where *B*_*c*_ is the maintenance power expenditure per cell. The change in the cellular dynamics (*dN/dt*) is subject to burst-like fluctuations arising from processes such as cell turnover, stochastic biochemical reactions and synchronized metabolic activity. This burst-like processes can be described by the Laplace distribution, which is the sum if two exponential processes [20]. We represent the aggregate cellular burst during one observation interval by

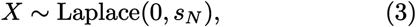

with size-dependent scale parameter

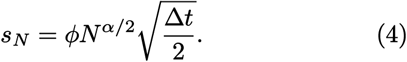

Here, *ϕ* controls the overall magnitude of stochastic fluctuations and *α* determines how their variance scales with organism size. Because Var 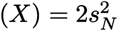, this formulation implies that the variance of the cellular fluctuations scale with the system size

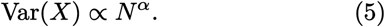

Independent cellular fluctuations correspond to *α* = 1, for which variances add linearly with cell number. Fully synchronized cellular activity corresponds to *α* = 2, whereas 1 *< α <* 2 represents partially correlated cellular dynamics. We assume that burst-like cellular activity introduces an additional metabolic cost proportional to the absolute burst magnitude. The total energy budget of the metabolism is therefore

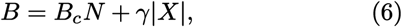

where *γ* converts cellular burst magnitude into metabolic expenditure. Note that at adult stage we do not account for growth component of metabolism. The parameter *γ* combines the energetic costs associated with cell turnover and stochastic cellular activity. Its mechanistic derivation from the full ontogenetic model is provided in the Supplementary Information.

### B. Expected metabolic rate and allometric exponent

For *X ~*Laplace(0, *s*_*N*_), the expected absolute burst magnitude is *E*[|*X*|] = *s*_*N*_. The expected adult metabolic rate is therefore

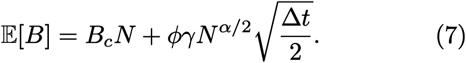

Since body mass is proportional to cell number, *m* = *m*_*c*_*N*, we define the local allometric exponent as

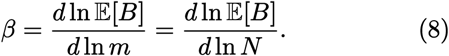

Differentiation gives

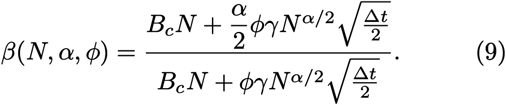

For 0 *<* α *<* 2, the effective exponent is bounded by α*/*2 *< β <* 1, with the precise value determined by the relative contribution of maintenance and stochastic expenditure. We fix the time step Δ*t* = 0.1 throughout the analysis. The resulting exponent reflects the relative contribution of linear cellular maintenance and size-dependent stochastic expenditure. When maintenance dominates, *β* → 1. When stochastic expenditure dominates, *β* → α*/*2.

### C. Simulation of metabolic fluctuations

For each organism size and noise parameter combination, two independent burst realizations were sampled,

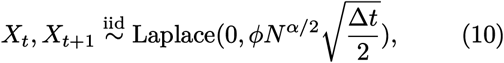

asand the corresponding metabolic rates were calculated

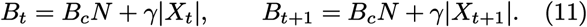

Metabolic fluctuations were quantified using the logarithmic return

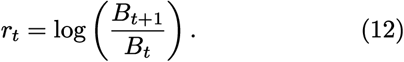

We evaluated α *∈* [1, 2], *ϕ ∈* [10^*−*6^, 10^*−*2^] and *N ∈* [10^10^, 10^16^]. For each parameter combination, we generated *n*_sim_ = 10^5^ independent pairs of metabolic observations.

### D. Information-theoretic analysis

In order to find the optimal noise structure and the corresponding allometric exponent, first we measure for each α the Kullback-Leibler (KL) divergence between the conditional fluctuation distribution of the mesoscopic fluctuations *r* and α and the pooled distribution of *r* across α:

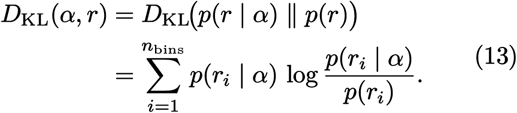

Here *p*(*r* |α) denotes the distribution of metabolic fluctuations generated under a given microscopic correlation exponent *α*, while *p*(*r*) represents the pooled distribution across all correlation exponents. The KL divergence therefore quantifies how strongly macroscopic fluctuations depend on the microscopic correlation structure. For each species size (i.e., each *N*), we evaluate *D*_KL_(α, *r*) across all α values between [1, 2]. Here, we define the information-neutral regime as the microscopic correlation structure for which the conditional macroscopic fluctuation distribution is least distinguishable from the pooled distribution. We interpret this condition as maximal robustness of the macroscopic fluctuation statistics to variation in microscopic correlation structure. The optimization proceeds in two stages. First, we identify the correlation exponent for which the macroscopic fluctuation distribution is least distinguishable from the pooled distribution across microscopic correlation structures. We refer to this minimum as the information-neutral regime. Second, we determine the fluctuation amplitude for which the corresponding information-neutral correlation exponent *α*_min_ is maximal across species sizes. This identifies the strongest microscopic correlation structure compatible with information neutrality (Fig. 1B).

**Figure 1.**
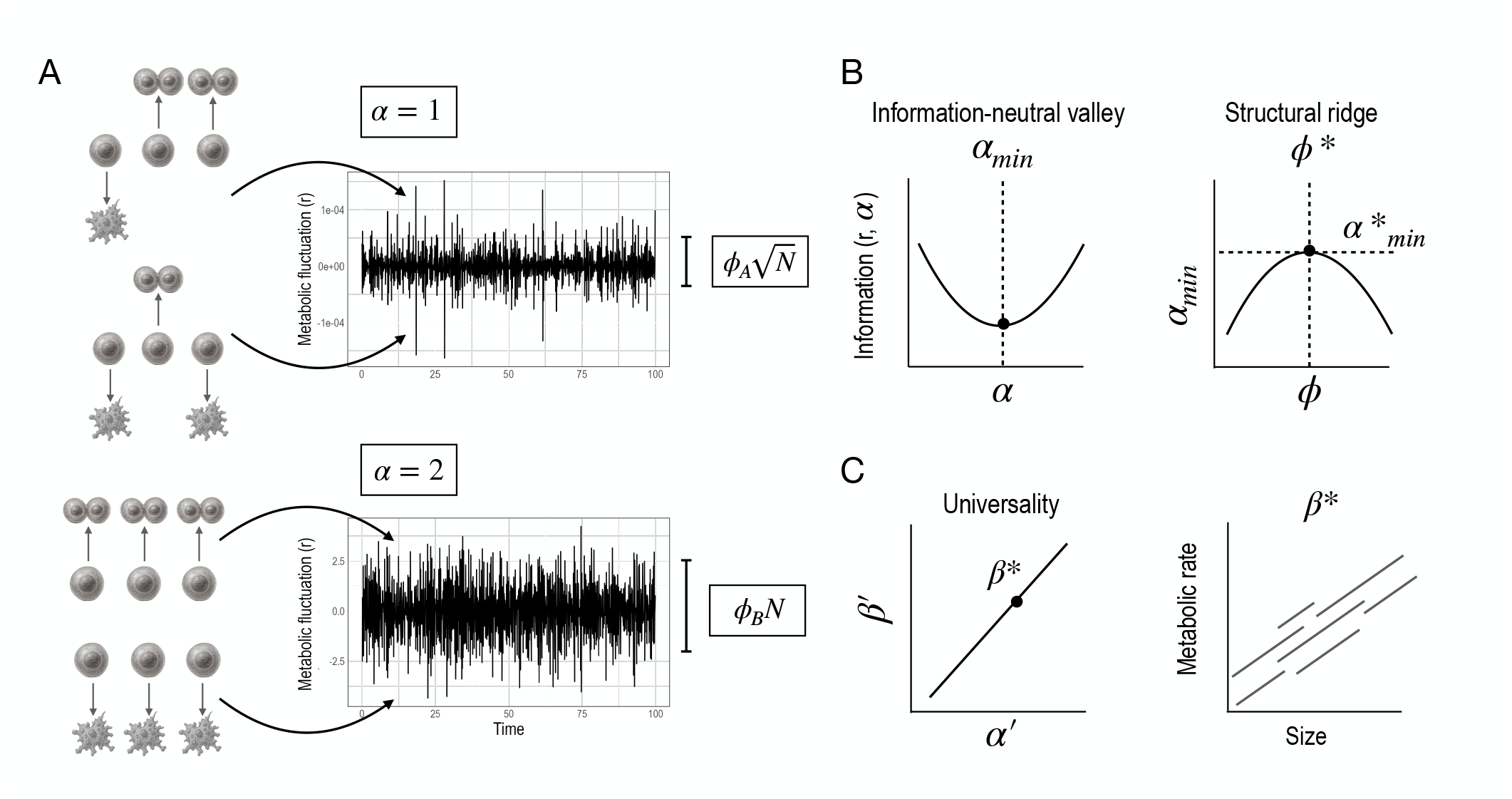
Conceptual framework of the information-neutral scaling mechanism. (A) Schematic illustration of metabolic fluctuations generated by stochastic cellular bursts. Uncorrelated cellular dynamics (top) produce weakly structured fluctuations, whereas correlated bursts (bottom) generate larger collective fluctuations at the organism level. (B) Identification of the information-neutral correlation exponent. For each value of the microscopic correlation exponent α, the distribution of metabolic fluctuations *p*(*r* |α) is compared with the pooled distribution *p*(*r*) using the KL divergence *D*_KL_(α, *r*). The minimum of the KL divergence identifies the information-neutral exponent α_min_, where the conditional macroscopic fluctuation distribution is least distinguishable from the pooled distribution across microscopic correlation structures. Second, we determine the optimal noise scale. The information-neutral exponent *α*_min_ is evaluated across different fluctuation amplitudes *ϕ*. The maximum of α_min_(*ϕ*) defines the structural ridge corresponding to the optimal fluctuation scale *ϕ*\*, which produces a universal metabolic scaling exponent *β*\* across species sizes. (C) Then using the identified optimal noise structure, we calculate the corresponding scaling exponent 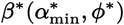.

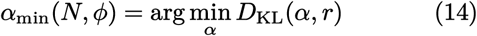

At the information neutrality point, α has minimal effect on the distribution of *r*. We define the information-neutral valley as the region in parameter space where the distribution of macroscopic fluctuations is least sensitive to variations in the microscopic correlation exponent α, quantified by a minimum in the *D*_*KL*_(α, *r*). This minimum defines the most robust correlation exponent α _*min*_. To reduce Monte Carlo noise, we smoothed (α, *D*_KL_) trajectories before locating the minimum. Then, we calculate the corresponding allometric exponents *β*(α _*min*_) for each body size across all *ϕ* values.

The goal is to determine whether there exists a species–independent optimal noise scale *ϕ*\* at which the allometric scaling exponent is universal. To do so, we calculate the overlap of the species-specific *α*_*min*_− *ϕ* curve representing the universal crossover. This value or interval represent the point where all species exhibit maximal robustness to microscopic noise and at the same time maximally correlated microscopic noise structure. For each species we identify the noise values *ϕ* at which robustness is maximal, i.e. when *α*_min_(*ϕ, N*) achieves its maximum over all possible *ϕ* values:

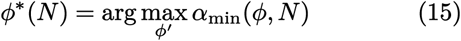

which defines the structural ridge or crossover region. Then, for each *ϕ*\*, we compute the corresponding allometric scaling exponent *β*\*.

To compare the structure of the species-specific trajectories independently of their absolute parameter ranges, we apply a min–max normalization,

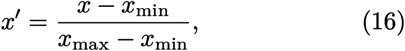

performed independently for each species size *N*. This transformation maps the trajectories onto a common dimensionless coordinate system and allows us to test whether their shapes collapse across organism sizes. Such trajectory collapse provides a phenomenological signature of universality. As an independent test of scale robustness, we examine whether the information-neutral minimum persists under temporal blocking of the fluctuation observable (see Supplementary Information).

As a complementary analysis, we quantify the sensitivity of the mesoscopic observable *r* to the scaling exponent *α* using the Fisher Information:

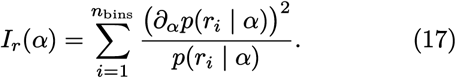

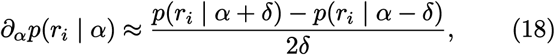

where is set to δ = 0.02. Then, calculating *I*_*r*_(α_*min*_) gives the sensitivity of the most robust correlation exponent to microscopic fluctuations.

## III. RESULTS

Our results reveal how information mapping from microscopic bursty cell dynamics to organism-level fluctuations shapes the species-level allometric exponent. First, we computed the KL divergence between the *α* and the organism-level metabolic fluctuations *r* across a α - *ϕ* grid for each species sizes (Fig.2A). In each parameter combination, a correlation exponent *α* _*min*_ can be identified, where the KL divergence reached its minimum (information-neutral valley). These minimum correlation exponents are in the range of *~* 1.4–1.8 across species sizes. At *α* _*min*_ the metabolic fluctuations *r*(*t*) retain the least sensitivity to the microscopic parameter *α*, indicating that the coarse-grained description is maximally insensitive to microscopic details.

To test for universality, we compared the normalized trajectories of *β*′(*α*_min_) and 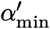 across organism sizes. The trajectories closely collapse onto a common relationship (Fig. 2B), indicating that the mapping between the information-neutral microscopic correlation structure and the macroscopic scaling exponent becomes approximately independent of organism size after normalization. This trajectory collapse therefore provides an information-theoretic signature of universality across the considered range of organism sizes.

**Figure 2.**
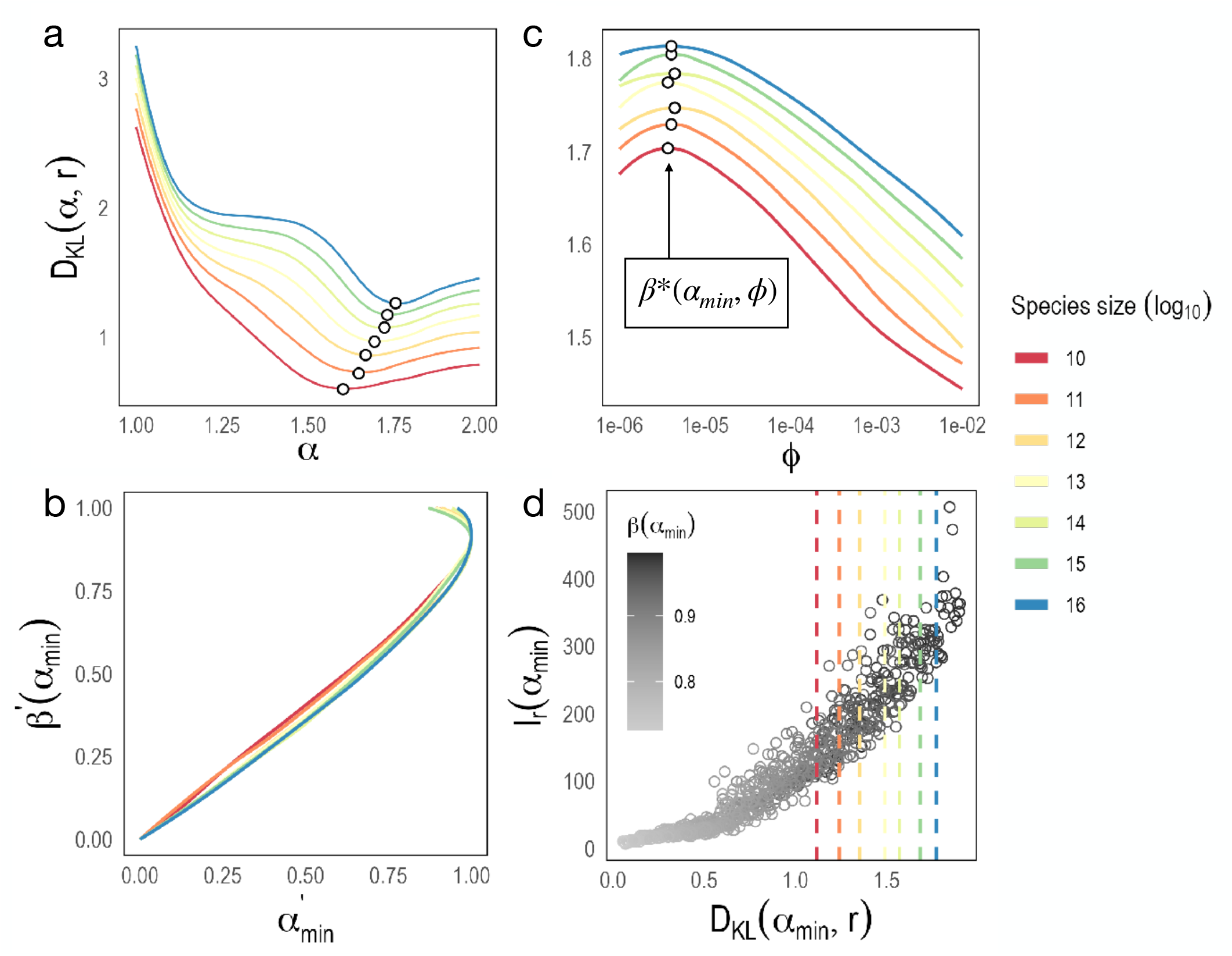
Sensitivity of stochastic metabolic fluctuations across species sizes. (A) An example of the KL divergence *D*_KL_(*α, r*) as a function of the microscopic correlation exponent *α* for a fixed fluctuation amplitude (log_10_(*ϕ*) = −4.5). The minimum identifies the information-neutral exponent *α* _min_ (black circles), where macroscopic metabolic fluctuations are least sensitive to the microscopic correlation structure. (B) Trajectories of the normalized scaling exponents *β*′(*α* _min_) and *α* ′ across different noise amplitudes *ϕ*. Each curve corresponds to a species size, and the collapse of trajectories indicates universal behavior. (C) The maximized *α* _min_(*ϕ*) identifies the structural ridge (black circles) corresponding to the optimal fluctuation scale *ϕ*\*, where microscopic correlations are strongest while remaining information-neutral. The corresponding metabolic scaling exponent *β*\* is shown. (D) Sensitivity of macroscopic fluctuations to the correlation exponent measured by the Fisher information.

As a next step, we identified the crossover region where the information-neutral correlation exponent *α*_min_ reaches its maximum as a function of the fluctuation amplitude *ϕ* (Fig. 2C). This defines the structural ridge *ϕ*\*, corresponding to the strongest microscopic correlation structure compatible with information neutrality. This minimax criterion identifies the most robust noise structure. Across six orders of magnitude in cell number, the corresponding scaling exponent is tightly constrained around *β*\* = 0.975 with a standard deviation of 0.004, despite a systematic shift in the microscopic correlation exponent from *α*\* = 1.70 to 1.81. At the structural ridge, stochastic expenditure contributes approximately 17% - 24% of the total expected metabolic expenditure, demonstrating that the near-linear scaling does not simply arise from the dominance of the deterministic maintenance term. Thus, distinct microscopic correlation structures and a substantial stochastic contribution converge onto a nearly size-independent macroscopic scaling exponent. Furthermore, we analysed the sensitivity of *r* with respect to *α* quantified by the Fisher Information. At the information-neutral states defined by the minima of *D*_KL_[*α*], Fisher information remains finite and does not coincide with its maximum (Fig.2D). Thus, information neutrality is distinct from a state of maximal sensitivity to microscopic fluctuations. For large *ϕ*, stochastic expenditure increasingly dominates the metabolic budget, and the allometric exponent approaches its stochastic limit, *β* → *α/*2.

## IV. DISCUSSION

The presented analysis shows that universality can arise from a simple information-neutrality criterion without requiring a phase transition. Rather than occurring at a point of maximal sensitivity to microscopic structure, universal metabolic scaling emerges in a non-critical crossover regime where organism-level fluctuation statistics are least distinguishable across microscopic correlation structures. In this regime, Fisher information remains finite and of intermediate magnitude, while species of different sizes converge toward a common metabolic scaling exponent. The theoretical optimum is consistent with recent evidence for near-linear metabolic scaling (*β* = 0.95) across the tree of life [17].

The universality observed here arises from the progressive loss of sensitivity to microscopic stochastic structure. Cellular bursts with different correlation properties generate distinct metabolic fluctuation statistics, yet within the information-neutral regime these differences have minimal influence on the macroscopic description. At the same time, the normalized relationship between microscopic correlation structure and the metabolic scaling exponent collapses across organism sizes, while the information-theoretic minimum remains stable under temporal aggregation. Together, these results indicate that robust macroscopic scaling can emerge from stochastic microscopic dynamics without requiring criticality or fine-tuning to a phase transition.

Such noncritical universal behaviors are increasingly recognized in biology. They appear in developmental scaling [23], neural population dynamics [28], biochemical reaction networks, protein aggregation, and scaling laws in ecosystems [2]. In these systems, robustness and universality emerge not from being near a phase transition, but rather from the natural elimination of microscopic detail as biological units are aggregated into larger scales [25]. These findings are similar to a previous model, where universal metabolic scaling arises from life-history optimisation [43].

The large variance in observed empirical diversity is more likely a result of a locally optimised trait space rather than global optimisation. Evolution can be formalised as a stochastic adaptive walk on rugged landscapes [18, 24] constrained by historical contingencies and developmental architecture [1, 13, 38], which prevent natural selection from globally optimizing traits. The average scaling exponent of *~*0.97 recovered in this analysis therefore represents a purely theoretical optimum. Note that a local minimum appears around *α* ≈ 1, which corresponds to uncorrelated noise structure.

This study offers a new theoretical perspective on how information neutrality can give rise to universal scaling laws. More broadly, the framework developed here suggests that information neutrality principles may underlie the universal scaling behavior of noisy processes. This model is generalizable to other complex systems.

